# Algorithm for Extraction of Sub-Structure from Co-Crystallized PDB Ligand for Selective Targeting

**DOI:** 10.1101/2020.02.02.931436

**Authors:** Om Prakash

## Abstract

An algorithm has been introduced for extraction of sub-structure from co-crystallized ligand of complex PDB file. Algorithm utilized location of atoms in 3D domain of complex of ligand & protein. It processed relative positioning of atoms for demarcation of influential part of ligand, which can provide potency to that ligand for binding with specific binding pocket of protein. Algorithm was validated with ligands co-crystallized with enzymes of different classes. Extracted sub-structures were validated (via evidence from database & literature) for selectivity towards one or more targets of same family.

## INTRODUCTION

Selective targeting of proteins by ligand in cellular processes is one of the major challenges in the area of toxicology and novel drug discovery. Solution for this challenge is looked into the structure of ligands or chemicals which are known with various cell signaling proteins. Many methods have been explained for this purpose as: Structural Alerts and Sub-Structure. These pre-existing methods used mechanism of structure comparison based on structural similarity, fingerprinting, and QSAR modeling etc. for identification of sub-structure or structural alerts. In previous studies, structural comparison is reported for identification of substructures. As European REACH works on the concept of weight-of-evidence approach for identification of bio-accumulative compounds. Quantitative Structure Activity Relationship (QSAR) based identification of structural alerts is used as one of factors in weight-of-evidence by REACH. It is to notify that here QSARs were used for extraction of structural alerts from compounds (Valsecchi et al., 2019). In another study, structural features based statistical QSAR models were defined structural alerts to flag potential chemical hazards (Alves et al., 2016). Molecular fragments with high chemical reactivity were also considered as structural alerts and were avoided to reduce toxicity in pharmaceuticals (Limban et al., 2018). Structure-metabolism studies are known to resolve reactive metabolite-related liabilities by "avoidance" strategies for exclusion of structural alerts and possible termination of reactive metabolite-positive compounds (Kalgutkar, 2019). In a study, to define the structural alerts for generation of reactive metabolites from drugs, systematic approach was used by involving macromolecules leading to immune responses (Claesson and Minidis, 2018)(Kalgutkar, 2019). In another study, to analyze the drug effect, classifier based approach was adopted for identification of association between drug substructures and protein domain. Here biologically relevant chemogenomic features were used in classifier for extraction of substructures (Tabei et al., 2012). In a study for predicting ligand-protein interaction, information of binding site was used along with sub-structure of ligand. To extract substructure, physical-chemical properties of binding site based approach was used (Wang et al., 2015). In a genome-wide screening of drug-target interaction, without requirement of 3D structure of target protein, sparse canonical correspondence analysis was performed by analyzing profiles of drug and targets simultaneously to extract sets of chemical substructures. These substructures were further exploited in a drug discovery process (Yamanishi et al., 2011).

Beyond existing methods, without using any structural comparative studies from a set of known ligands, X-ray crystallographic complex structure can be used directly for extraction of potentially influential sub-structure(s) from ligand. Here it is assumed that high selectivity information is hidden in location of atoms in 3D domain of complex of ligand & protein. Therefore mechanical transformation of relative positioning can be used to gather information for demarcating influential part of ligand within the complex system of specific binding pocket of protein. Therefore, a generalized algorithm has been introduced for extraction of sub-structure directly from co-crystallized protein-ligand complex PDB file.

## MATERIALS AND METHOD

Algorithm has been developed and implemented for extraction of sub-structure of ligand which can be potentially involved in molecular interaction with specific protein. Assumptions were made for hypothesizing algorithm as: (i) each atom has same mechanical capability to affect other atom; (ii) co-crystal structure is at most optimized stable/stagnant state; (iii) during ligand-protein interaction, the co-crystallized ligand has found stable interaction conformation; and (iv) theoretically it was also assumed that dynamicity of protein and ligand, ultimately achieve a combination where ligand get pegged at one or multiple locations inside the force field of protein domain. Each pegging hook is headed by single atom (here called ‘leader atom’). Leader atom is followed by other atoms with decreasing stretch. Impact of hook decreases with distance from leader atom and which ultimately converses into zero. Selectivity of ligand towards target is presented by the most stretched hook atom. Ligand atom closest to protein was considered as the most efficient atom to pull other ligand atoms towards protein. This selection of leader atom as well as defining the stretch gradient score have been implemented into algorithm named ‘Relative Unified Mechanical Skill Score of Atom’ (RUMSSA). It extracts sub-structure from co-crystallized ligand for selective targeting. It process PDB file directly. It is a generalized as well as unbiased method to process co-crystallized PDB structure. This method ensures the molecule interaction, since picked from PDB itself.

### Algorithm

Relative Unified Mechanical Skill Score of Atom (RUMSSA) is a algorithm; which is capable to mark-out a sub-structure part of ligand from PDB file containing two molecules (Protein-Ligand) interacting each other. Calibration is required for each complex. Therefore program performance should be calibrated with customizing distance value. RUMSSA algorithm steps have been defined as: **Processing with whole molecule**: whole complex is assumed into 3D space. Location of each atom is traced into 3D space. Calculate distance between each pair of atoms. A sphere is assumed around each atom (radius of sphere is customizable, default 5Å). Each atom is initializing with RUMSSA score value of zero. **Processing with each atom**: Select atom pairs within the sphere (i.e. has inter-atomic distance of >0 & <=radius of sphere). Update local RUMSSA value of atom as (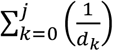) of Ligand within sphere around it. Repeat this process for each ligand atom. Sort the atoms on the basis of initial RUMSSA value. Atom with RUMSSA value of ‘Zero’, will be named as ‘Leader atom’. Calculate distance of each Ligand-atom from ‘Leader atom’, within a customized range (default 5Å) and sort them. Now calculate ‘Relative RUMSSA i.e. R_Di_’ in relation of ‘RUMSSA of Leader atom’ and distance (i.e. Di) of each atom from ‘leader atom’. Calculate ‘max of R_Di_ vector’: this will be ‘Threshold’ for demarcating atoms starting from leader atom. Global RUMSSA value for each atom of ligand can be defined as following:

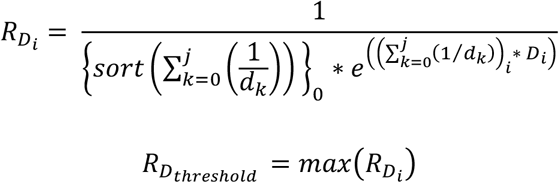

Condition for picking out the sub-structure selective for target protein: Atom should have Global RUMSSA value < R_DThreshold_ and Local RUMSSA value of atom should be common of sorted & non-sorted atom.

### Demonstration of algorithm

Implementation of algorithm was described with PDB: 3EQM containing complex of Aromatase along with Androstenedione.

### Approval of algorithm

Since RUMSSA processes co-crystallized PDB complex structures, therefore RUMSSA algorithm was validated with ligands co-crystallized with enzymes of different classes. Extracted sub-structures were proved for selectivity towards one or more targets of same family. Algorithm has been implemented for extraction of sub-structure(s) potentiating ligand for binding with specific protein. To perform generalized approval of algorithm, PDBs were selected from protein data bank with text Search for: drug and TAXONOMY is just Homo sapiens (human) and Chain Type: there is a Protein chain but not any DNA or RNA or Hybrid and Resolution is 1.499 or less and Stoichiometry in biological assembly: Stoichiometry is MONOMER and Resolution is between 0.6 and 1.099 Å. As result three groups of enzymes Oxidoreductases, Lyases and Hydrolases were considered for analysis.

## RESULTS AND DISCUSSION

Ligand molecule interacts with protein to perform some activity. But not whole molecule is conserve for performance against specific protein. One or more sub-structure bears some structural & conformational combination for potentiating the selectivity towards the protein molecule. Therefore to know and reuse the sub-structure, it becomes important to extract sub-structure from ligand. But problem is that the demarcation of sub-structure is not defined. To define method, theme of assumption has been re-evolved. Algorithm has been defined on basis of mechanical behavior of ligand atoms. Algorithm has been made in generalized scheme, so that it can be used for any ligand. Since ligand is already present in 3D field domain of protein, therefore atomic locations can be redefined for describing hypothesis of mechanical behavior for generalized scheme development. To define the mechanical behavior, reference point has been defined as leader atom. Theoretically, leader atom show highest ability to interact with protein and rest of ligand atoms follow the leader’s way of approaching protein. Therefore this theme of hypothesis was defined in terms of ‘Relative Unified Mechanical Skill Score of Atom’. It was assumed that atoms those follow the leader atom should be the part of potentiating sub-structure; and those don’t follow the leader atom should not be a part of substructure. Algorithm judges the complex molecule in two aspects: Global & Local. Global aspect covers whole molecule under observation, while local aspect observe each individual atom of ligand. Where the global & local aspects covers a common part, is marked as sub-structure required for selectivity.

Since algorithm considered complex molecule observation at global & local aspect, therefore it became important to calibrate the algorithm for accurate results. Although RUMSSA consider all possible leader atoms, but it provides most accurate sub-structure when it is calibrated for single leader atom. Since algorithm considers 3D complex PDB structure, therefore Location of each atom is observed into 3D space. Ligand atoms trapped into the force field of protein, unravel the hidden pattern for molecular selectivity. Therefore RUMSSA calculates distance between each pair of atoms. Since each atom of ligand is influenced by their neighbor atoms, therefore a range of area was assumed into sphere to achieved average relative influence on each atom. Each atom was assumed with unified RUMSSA value, which made the algorithm unbiased as well as generalized for any ligand molecule. Therefore this value can also be used for inter-molecule comparative study. RUMSSA represents a hypothetical parameter which depends only on distance of ligand atoms relative the leader atom optimized within the force-field of protein; therefore this mechanical transformation is observed with inter-atomic distance. This observation is bidirectional i.e. global as well as local. And both aspects of observations are independent to each other. Finally both observations are read through relative commonality to achieve the required sub-structure demarcation. Therefore sub-structure is picked on the basis of global relative RUMSSA value as well as Local RUMSSA value of atom.

To exemplify the working strategy of RUMSSA, demonstration of algorithm was done with PDB 3EQM, in which Ligand was totally covered with protein, and RUMSSA picked a substructure of most important section. Figure 1 described distribution of global RUMSSA value starting from leader atom. Highest RUMSSA value is threshold, beyond which ligand will be uninfluenced of leader atom.

**Figure 1.**
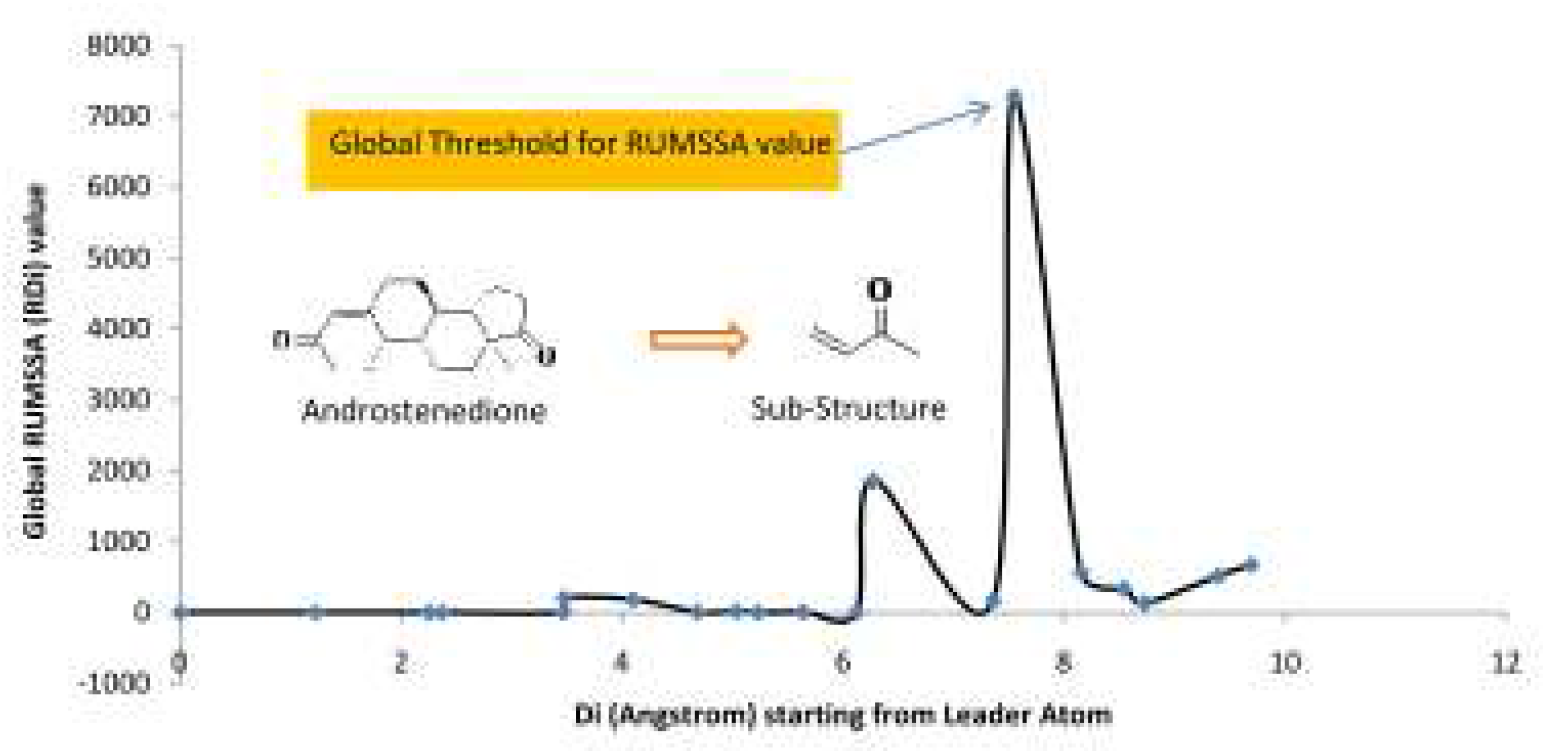
Demonstrating execution of RUMSSA algorithm for extraction of Sub-Structure from PDB: 3EQM containing Aromatase & Androstenedione

Furthermore, to provide the approval to algorithm, algorithm was evaluated with PDBs for Human protein with highest resolution range of 0.6 to 1.099Å. Purpose of these case studies as experimental validation of performance of algorithm was to check whether identified sub-structures through RUMSSA are also present in known ligands targeting specific protein. To perform generalized validation, protein data bank was filtered with 03 groups of enzymes Oxidoreductases, Lyases and Hydrolases and was considered for analysis. Identified sub-structures from known PDBs have been tabulated in Table 1 & 2. It was found that extracted sub-structures also existed in other known PDB of same protein or protein of same class. Enzyme isoforms were also found to be bounded with those sub-structures.

**Table 1.**
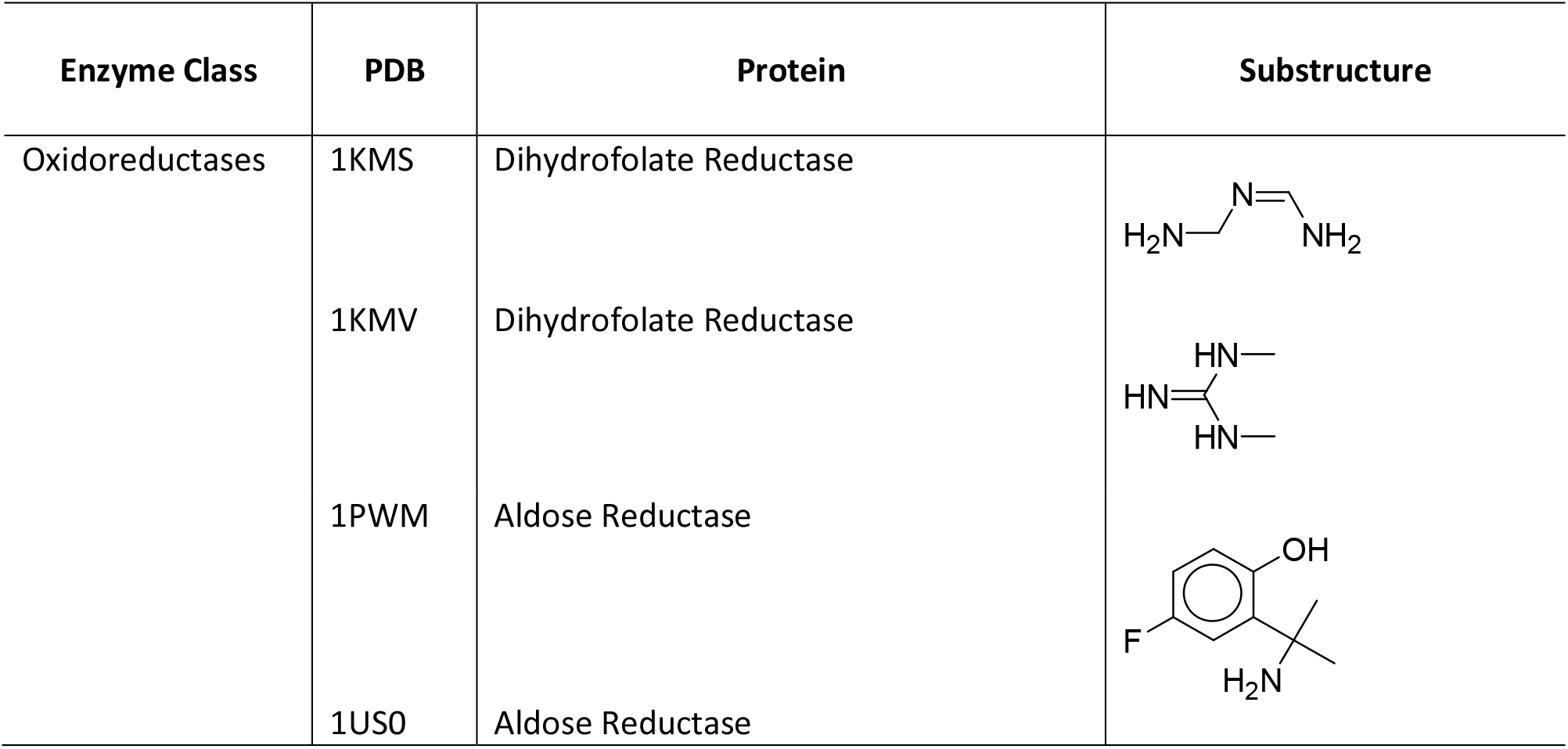
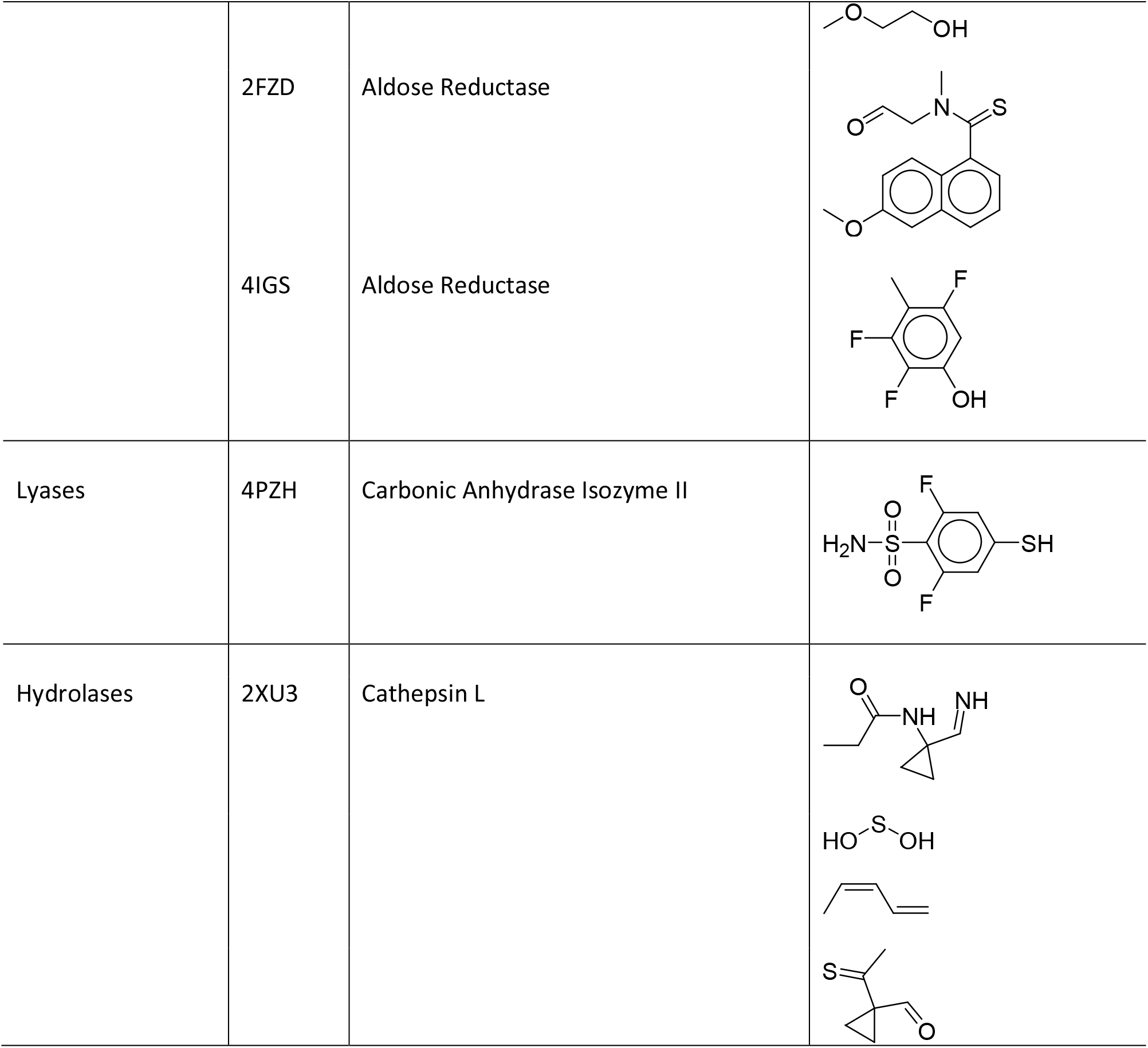
Identified Sub-Structures from known PDBs of three different classes of enzymes. Smaller sub-structures were found non-selective by covering a large population of compounds. This suggests that size of sub-structure may be a criterion for selectivity towards the target proteins.

**Table 2.**
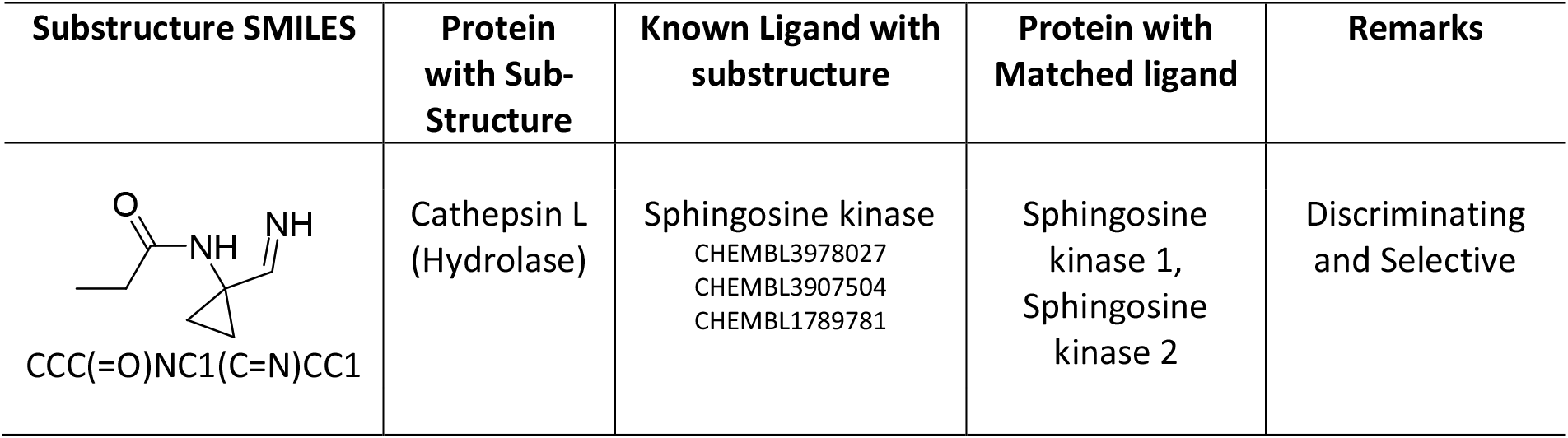
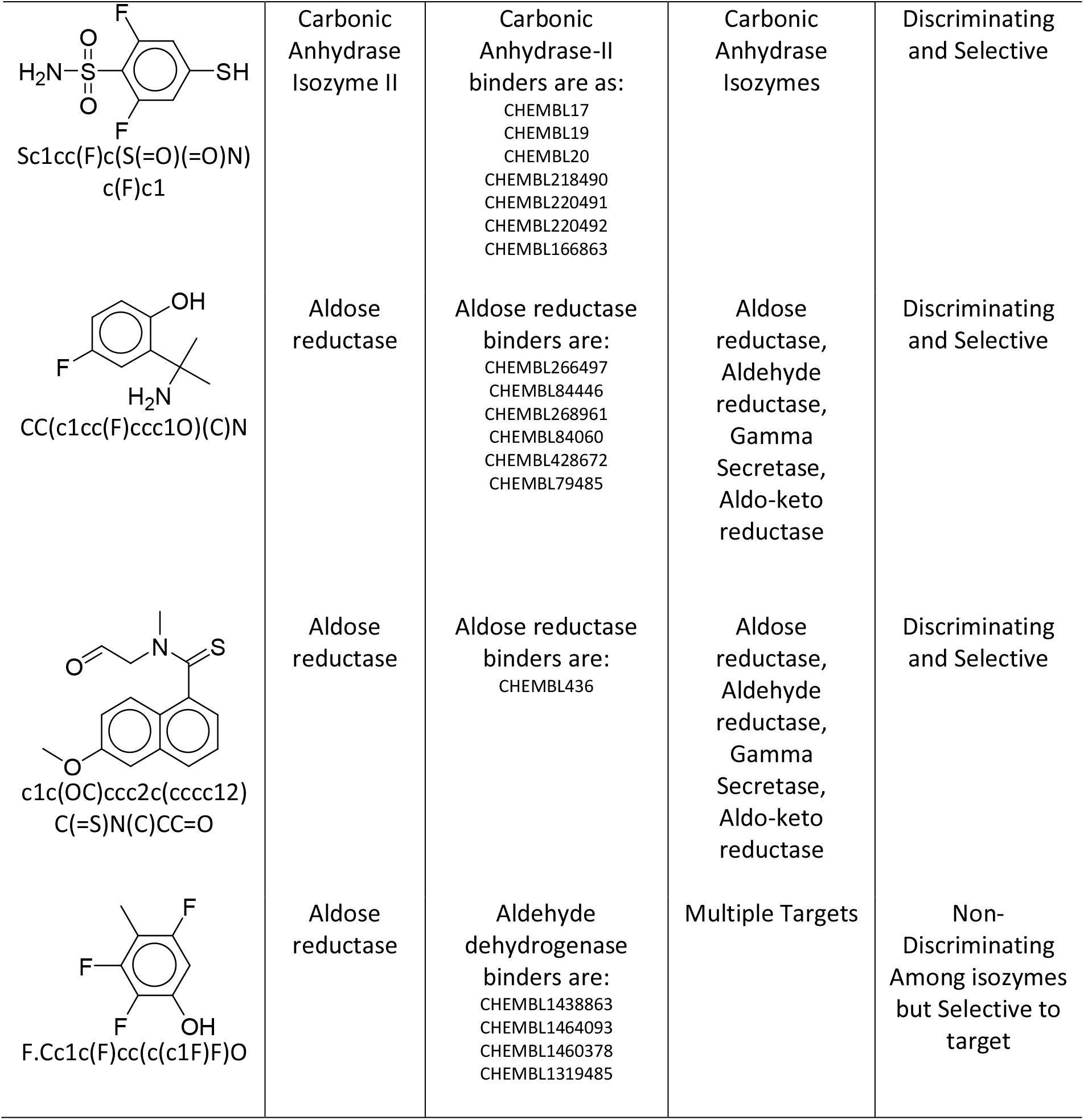
Sub-Structure SMILES vs. Known ligands with Protein classes, other known ligands, proteins and remark. From Table 1, larger sub-structures showed selectivity towards target so included in this table.

| Substructure SMILES | Protein with Sub-Structure | Known Ligand with substructure | Protein with Matched ligand | Remarks |
| --- | --- | --- | --- | --- |
| CCC(=O)NC1(C=N)CC1 | Cathepsin L (Hydrolase) | Sphingosine kinase<br>CHEMBL3978027<br>CHEMBL3907504<br>CHEMBL1789781 | Sphingosine kinase 1,<br>Sphingosine kinase 2 | Discriminating and Selective |
| Sc1cc(F)c(S(=O)(=O)N)c(F)c1 | Carbonic Anhydrase Isozyme II | Carbonic Anhydrase-II binders are as:<br>CHEMBL17<br>CHEMBL19<br>CHEMBL20<br>CHEMBL218490<br>CHEMBL220491<br>CHEMBL220492<br>CHEMBL166863 | Carbonic Anhydrase Isozymes | Discriminating and Selective |
| CC(c1cc(F)ccc1O)(C)N | Aldose reductase | Aldose reductase binders are:<br>CHEMBL266497<br>CHEMBL84446<br>CHEMBL268961<br>CHEMBL84060<br>CHEMBL428672<br>CHEMBL79485 | Aldose reductase, Aldehyde reductase, Gamma Secretase, Aldo-keto reductase | Discriminating and Selective |
| c1c(OC)ccc2c(cccc12)C(=S)N(C)CC=O | Aldose reductase | Aldose reductase binders are:<br>CHEMBL436 | Aldose reductase, Aldehyde reductase, Gamma Secretase, Aldo-keto reductase | Discriminating and Selective |
| F.Cc1c(F)cc(c(c1F)F)O | Aldose reductase | Aldehyde dehydrogenase binders are:<br>CHEMBL1438863<br>CHEMBL1464093<br>CHEMBL1460378<br>CHEMBL1319485 | Multiple Targets | Non-Discriminating Among isozymes but Selective to target |

As done in previous studies, structural comparative studies have drawbacks of assurance of existence of sub-structure. Reason behind this is non-confidant collection of initial dataset for development of models. As identification of sub-structure based on QSAR models, with set of positive & negative sets of dataset, does not assure about molecular interaction as well as selectivity towards target (Valsecchi et al., 2019). In another study, structural features were used, which does not assure that relevant fingerprint as required for target selectivity (Alves et al., 2016). Use of molecular fragments does not assure about coverage of inter-fragment connection based substructures (Limban et al., 2018). Similarly there are other aspects which are not covered by ligand based comparative studied with set of compounds. Beyond these, RUMSSA read ligand within the force-field of protein; therefore it is unbiased to the set of ligands. Ligand based comparative modeling does not use conformation information of ligand, while RUMSSA uses location of each atom which is the subject of molecular conformation. It must be noted that RUMSSA will be implemented with 3D structure complex structure of Ligand-Protein. During analysis of case studies it was observed that small sub-structure did not showed selectivity towards target, which larger sub-structures did (Table 1, 2). Therefore it has been marked that, may be the size of sub-structure as factor for selectivity towards target. The extracted sub-structures will be of use as chemical alerts for database searching as well as development of potential lead molecules for various biological activities.

### Availability of script

Algorithm script can be accessed through author.

## CONCLUSION

RUMSSA was found to be efficient in extraction of selective sub-structures from PDB file of co-crystallized structure. It was found that size of sub-structure may be a factor for selective targeting. This sub-structures extraction algorithm will be useful in novel drug discovery and toxicological research.

## ACKNOWLEDGEMENT

Author express gratitude to *The Institute of Mathematical Sciences*, Chennai-600113, India for providing research facilities as well as DAE Post-Doctoral Fellowship (PDF 214). Author is also thankful to Dr. Amit Singh (PDF Mathematics, IMSc) for mathematical proof editing.

## REFERENCES

Alves, V.M., Muratov, E.N., Capuzzi, S.J., Politi, R., Low, Y., Braga, R.C., Zakharov, A. V., Sedykh, A., Mokshyna, E., Farag, S., Andrade, C.H., Kuz’Min, V.E., Fourches, D., Tropsha, A., 2016. Alarms about structural alerts. Green Chem. https://doi.org/10.1039/c6gc01492e

Claesson, A., Minidis, A., 2018. Systematic Approach to Organizing Structural Alerts for Reactive Metabolite Formation from Potential Drugs. Chem. Res. Toxicol. https://doi.org/10.1021/acs.chemrestox.8b00046

Kalgutkar, A.S., 2019. Designing around Structural Alerts in Drug Discovery. J. Med. Chem. https://doi.org/10.1021/acs.jmedchem.9b00917

Limban, C., Nuţă, D.C., Chiriţă, C., Negreş, S., Arsene, A.L., Goumenou, M., Karakitsios, S.P., Tsatsakis, A.M., Sarigiannis, D.A., 2018. The use of structural alerts to avoid the toxicity of pharmaceuticals. Toxicol. Reports. https://doi.org/10.1016/j.toxrep.2018.08.017

Tabei, Y., Pauwels, E., Stoven, V., Takemoto, K., Yamanishi, Y., 2012. Identification of chemogenomic features from drug-target interaction networks using interpretable classifiers. Bioinformatics. https://doi.org/10.1093/bioinformatics/bts412

Valsecchi, C., Grisoni, F., Consonni, V., Ballabio, D., 2019. Structural alerts for the identification of bioaccumulative compounds. Integr. Environ. Assess. Manag. https://doi.org/10.1002/ieam.4085

Wang, C., Liu, J., Luo, F., Deng, Z., Hu, Q.N., 2015. Predicting target-ligand interactions using protein ligand-binding site and ligand substructures. BMC Syst. Biol. https://doi.org/10.1186/1752-0509-9-S1-S2

Yamanishi, Y., Pauwels, E., Saigo, H., Stoven, V., 2011. Extracting sets of chemical substructures and protein domains governing drug-target interactions. J. Chem. Inf. Model. https://doi.org/10.1021/ci100476q

